# A pitfall for machine learning methods aiming to predict across cell types

**DOI:** 10.1101/512434

**Authors:** Jacob Schreiber, Ritambhara Singh, Jeffrey Bilmes, William Stafford Noble

## Abstract

Machine learning models used to predict phenomena such as gene expression, enhancer activity, transcription factor binding, or chromatin conformation are most useful when they can generalize to make accurate predictions across cell types. In this situation, a natural strategy is to train the model on experimental data from some cell types and evaluate performance on one or more held-out cell types. In this work, we show that when the training set contains examples derived from the same genomic loci across multiple cell types, the resulting model can be susceptible to a particular form of bias related to memorizing the average activity associated with each genomic locus. Consequently, the trained model may appear to perform well when evaluated on the genomic loci that it was trained on but tends to perform poorly on loci that it was not trained on. We demonstrate this phenomenon by using epigenomic measurements and nucleotide sequence to predict gene expression and chromatin domain boundaries, and we suggest methods to diagnose and avoid the pitfall. We anticipate that, as more data and computing resources become available, future projects will increasingly risk suffering from this issue.

---

Machine learning has been applied to a variety of genomic prediction problems, such as predicting transcription factor binding, identifying active cis-regulatory elements, constructing gene regulatory networks, and predicting the effects of single nucleotide polymorphisms. The inputs to these models typically include some combination of nucleotide sequence and signals from epigenomics assays.

Given such data, the most common approach to evaluating predictive models is a “cross-chromosomal” strategy, which involves training a separate model for each cell type and partitioning genomic loci into some number of folds for cross-validation (Figure 1a). Typically, the genomic loci are split by chromosome. This strategy has been employed for models that predict gene expression [1, 2, 3], elements of chromatin architecture [4, 5], transcription factor binding [6, 7], and cis-regulatory elements [8, 9, 10, 11, 12, 13]. Although the cross-chromosomal approach measures how well the model generalizes to new genomic loci, it does not measure how well the model generalizes to new cell types. As such, the cross-chromosomal approach is typically used when the primary goal is to obtain biological insights from the trained model.

**Figure 1:**
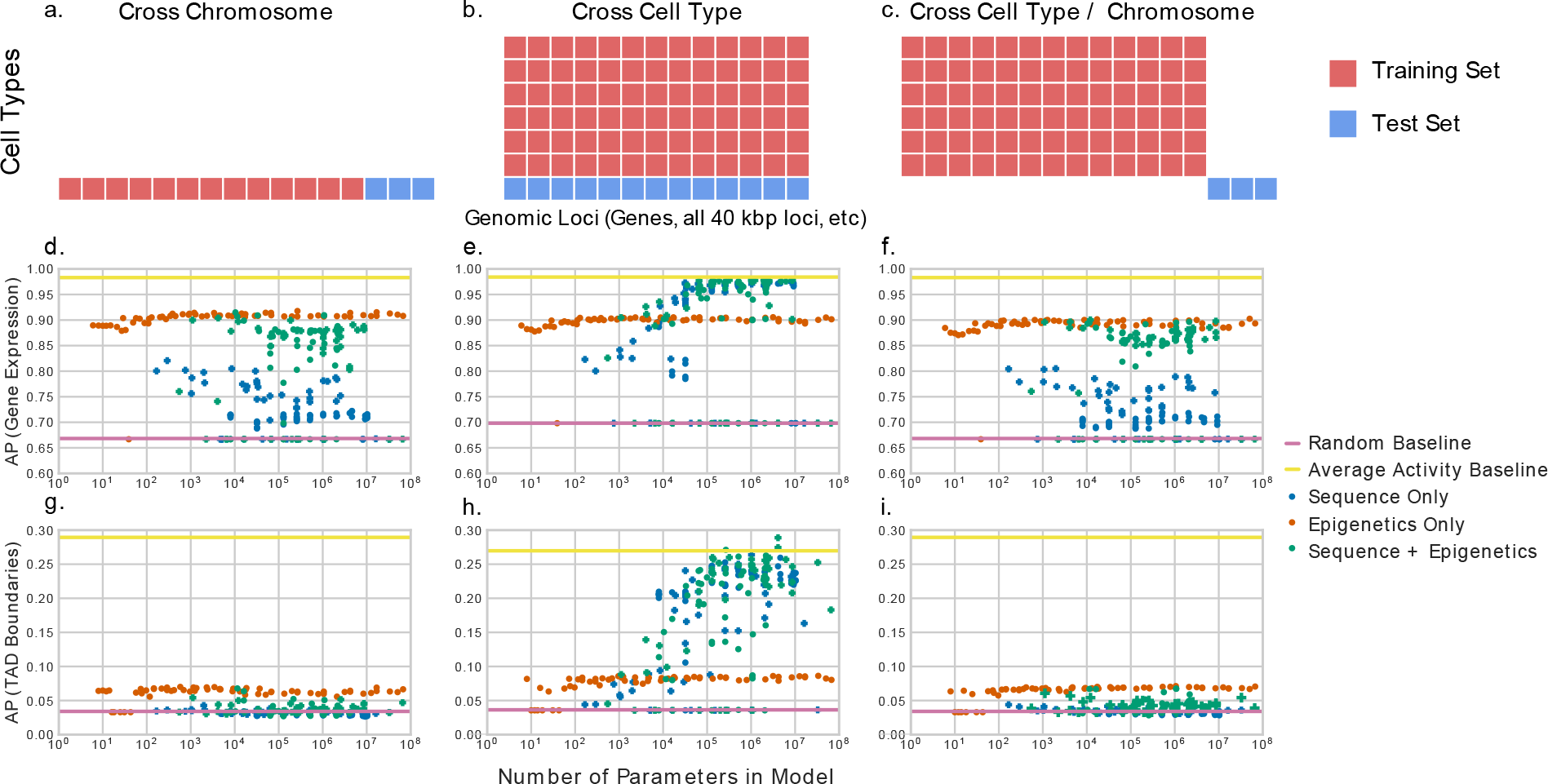
The performance of neural network models of varying complexity in three predictive settings on two tasks. Schematic diagrams of (a) cross-chromosome, (b) cross-cell type, and (c) hybrid cross-cell type / cross-chromosomal model evaluation schemes. (d–f) The figure plots the average precision (AP) of a machine learning model predicting gene expression as a function of model complexity. Evaluation is performed via (d) cross-chromosome, (e) cross-cell type, and (f) a combination of cross-chromosome and cross-cell type validation. In each panel, each point represents the test set performance of a single trained model. (g–i) is the same as (d–f) but predicting TAD boundaries rather than gene expression.

An alternative, “cross-cell type” validation approach can be used to measure how well a model generalizes to a new cell type. This approach involves training a model in one or more cell types and then evaluating it in one or more other cell types (Figure 1b). Researchers have used this approach to identify cis-regulatory elements [14, 15, 16, 17, 18], impute epigenomics assays that have not yet been experimentally peformed [19, 20], and predict CpG methylation [21]. The cross-cell type strategy is typically adopted when the goal is to yield predictions in cell types for which experimental data is not yet available.

In this work, we point out a potential pitfall associated with cross-cell type validation, in which this evaluation strategy leads to overly optimistic assessment of the model’s performance. In particular, we observed that models evaluated in a cross-cell type setting seem to perform better as the number of parameters in the model increases. To illustrate this phenomenon, we train a series of increasingly large neural networks to predict gene expression as measured by RNA-seq in the H1 cell line (E003), evaluating each model using the cross-chromosomal and the cross-cell type approaches. As input, each model receives a combination of nucleotide sequence and epigenomic signal from examples in the H1 cell line or 55 other cell lines, depending on evaluation setting (see Methods). In every case, we evaluate model performance using the average precision score relative to a binary gene expression label (“high” versus “low” expression). In the cross-chromosome setting, the performance of the models remains fairly constant as the complexity of the learned model increases (green points in Figure 1d). On the other hand, the cross-cell type results show a surprising trend: using more complex models appears to yield consistently better results, even as the models become very large indeed (up to 100 million parameters; Figure 1e).

To see that this apparently good predictive performance is misleading, we perform a third type of validation, a hybrid “cross-chromosome / cross-cell type” approach in which the model is evaluated on loci and cell types that were not present in the training set (Figure 1c). This approach eliminates the positive trend in model performance as a function of model complexity (Figure 1f). Very similar trends are seen when we train neural networks to predict the locations of topologically associating domain (TAD) boundaries in the H1 cell line (Figure 1g–1i). Further, these results do not appear to be specific to deep neural networks, as gradent boosted decision tree classifiers show similar trends as the number of trees increases (Supplementary Figure S1).

Interestingly, we note that the performance of models that use only epigenomic signal is fairly invariant to the number of parameters in the model. This suggests that there is an association between our representation of histone modification and gene expression that requires only few parameters to capture, such as H3K4me3 and H3K4me1 generally being activating marks and H3K27me3 generally being a repressive mark. Indeed, when we project the epigenomic signal into two dimensions, we observe regions in 2D where highly expressed genes can be easily separated from lowly expressed genes and regions where separation seems difficult by any method (Supplementary Figure S2a/b). We see a similar trend in model performance on synthetic Gaussian data when the two classes partially overlap (Supplementary Figure S3b). This is likely because while larger models have greater potential to overfit to samples in the overlap, the overall metric is not significantly influenced because the majority of points can be correctly classified by a simple rule.

The following two observations suggest that the positive trend in Figure 1e arises because more complex models effectively “memorize” the genomic location associated with expressed versus non-expressed genes. First, if we train a model using only the epigenomic signal, without including the nucleotide sequence as input, then the model performance no longer improves as a function of model complexity (orange points in Figure 1e); conversely, providing only nucleotide sequence as input yields very good performance across many cell types (blue points in Figure 1e). Second, comparison to a suitable baseline predictor—namely, the average expression value associated with a given locus across all cell types in the training set—outperforms any of the trained models (solid yellow line in Figure 1e). Thus, it seems that the more complex neural networks achieve good performance by effectively remembering which genes tend to exhibit high or low expression across cell types. Furthermore, though we demonstrate here that models may use nucleotide sequence to memorize gene activity, the phenomenon is more general, in the sense that any signal that is constant across cell types can be exploited in this fashion. Examples include features derived from the nucleotide sequence—k-mer counts, GC content, nucleotide motifs occurences, or conservation scores—or even epigenomic data when the input is signal from a constant set of many cell types rather than a single cell type.

It is worth pointing out that, from a machine learning perspective, the neural network is not doing anything wrong here. On the contrary, the neural network is simply taking advantage of the fact that most genomic or epigenomic phenomena that are subjected to machine learning prediction exhibit low variance, on average, across cell types. For example, the gene expression level of a particular gene in a particular cell type is much more similar, on average, to the level of that same gene in a different cell type than it is to the level of some other gene in the same cell type. Similarly, many transcription factors bind to similar sets of sites across cell types, most pairs of promoters and enhancers will never interact, and most regions of the genome are unlikely to ever serve as TAD boundaries.

This pitfall can be identified in several ways. First, comparison of model performance to an appropriate baseline, such as the average activity in the training cell types at the given locus (yellow lines in Figure 1e,f,h,i), will often show that an apparently good model underperforms this relatively simple competitor. As an example, we note that this average activity baseline outperforms two of the top four participants in the ENCODE-DREAM transcription factor binding challenge at predicting CTCF in the iPSC cell line when the models were evaluated on loci that they were also trained on (Supplementary Figure S4). This demonstrates that it is not sufficient to perform evaluations in the hybrid setting, but that one should also perform evaluations in the same setting that one would practically use the model. Importantly, even when a model is evaluated in the hybrid setting, one should compare to the performance of this baseline to demonstrate that a downstream user of the predictions would not be better suited by using this simple method. If the trained machine learning model cannot outperform this “average activity” baseline, then the predictions from this model may not be practically useful.

Second, the performance of the model can be more fully characterized by partitioning genomic loci into groups according to their variability across cell types and then evaluating model performance separately for each group (Supplementary Figure S5). This partitioning removes the predictive power of the average activity; thus, models that have memorized this average activity will no longer perform well. Indeed, we observe that models that use only nucleotide sequence appear to perform well in the cross-cell type setting but perform markedly worse when evaluated in this partitioned manner.

There are several approaches that may improve the cross-cell type predictive performance of models that underperform the average activity baseline. A natural approach is to use the average activity directly as a feature in the machine learning model. As an input feature, the average activity would have to be defined over a held-out subset of the training cell types to prevent a leakage of information between the input features and the target labels, but would nonetheless be a straightforward addition. Another approach would be to phrase the prediction problem not as predicting the activity directly, but predicting the difference from the average activity at that locus for that specific cell type. This approach allows the model to focus on learning cell type-specific differences. It is likely that different architectural decisions would need to be made when predicting the difference from the average activity instead of the activity directly, such as changing the loss function from a classification one to, potentially, a regression one. These strategies both explicitly use the average activity during the training procedure, with the first using it as an input and the second using it to create a new set of labels.

While most cross-cell type predictive tasks would benefit from a comparison to the average activity baseline, it is important to note in some settings beating the average activity baseline is not necessary. One such setting is when the goal is to inspect the trained model to derive new biological insights. For example, a researcher studying chromatin architecture may build a machine learning model that aims to predict genomic 3D structure using epigenomic state. If the trained model does not outperform using the average chromatin architecture from other cell types, then one may not be inclined to use the resulting predictions in downstream analyses. However, inspecting the model may still yield useful insights into the association between certain epigenomic marks and chromatin architecture. Another such setting is the semi-supervised setting, where only a portion of labels are known in advance and the goal is to identify previously unidentified annotations. In this case, because the full set of true labels is not known in advance, a comparison to the average activity may be a poor estimator of the ability of the model to identify novel elements. A final setting is that of anomaly detection, where one identifies regions that are poorly modeled for further study. In both of these settings, it is still informative to compare the performance of the models to the average activity baseline to demonstrate the strength of the predictive model.

As more data becomes available, we anticipate that more projects will risk suffering from the pitfall that we describe. Fortunately, avoiding this trap is straightforward: compare model performance to a baseline method that extracts the experimental signal from one or more training cell types. The simplest such strategy is to average the signal at a given locus across all training cell types. A more sophisticated strategy would be to use as a baseline the activity of a cell type in the training set whose biological activity, such as epigenomic state, is empirically similar to the target cell type. Regardless, comparing a model’s predictions to the activity of the training cell types is a necessary component of demonstrating the utility of the model.

## Methods

### Data sets

Nucleotide sequence are extracted from the hg19 reference genome. Before input to our models, each sequence is one-hot encoded such that each genomic position is represented by four bits, of which only a single one is 1. For the task of active gene prediction, a 2 kbp region is extracted upstream of the transcription start site, accounting for the strand of the gene. For the task of TAD boundary prediction, a 2 kbp region is extracted from the middle of the 40 kbp region to be considered.

The ChIP-seq, DNase-seq and gene expression RPKM values were downloaded from the Roadmap compendium (https://egg2.wustl.edu/roadmap/data/byFileType/signal/consolidated/macs2signal/pval/ and https://egg2.wustl.edu/roadmap/data/byDataType/rna/expression/). Each ChIP-seq and DNase-seq experiment is reported using – log_10_ p-values, indicating the statistical significance of the enrichment of the measured phenomenon at each genomic position. Additionally, these tracks are *arcsinh* transformed, which is similar to a log transform and is a standard technique to reduce the effect of outliers on the model. After this transformation, the average signal value for each epigenomic mark across the 2 kbp region of interest is used as input to our models. We used experimental measurements of H3K4me3, H3K27me3, H3K36me3, H3K9me3, and H4K3me1 for the prediction of gene expression, and additionally measurements of DNase-seq and H3K27ac for predicting TAD boundaries.

Gene bodies were defined as GENCODE v19 gene elements (https://www.gencodegenes.org/human/release_19.html) on chr1–22, resulting in 17,951 gene bodies for each of 56 different human cell types. We define active genes as those that have an RPKM value of *>* 0.5.

TAD boundary calls were obtained from the supplementary material of [22] for the seven cell lines TRO, H1, NPC, GM12878, MES, IMR90, and MSC. These calls are binary indicators and were specified at 40 kbp resolution.

Predictions from the top four participants in the ENCODE-DREAM challenge and the CTCF test set labels were provided by the ENCODE-DREAM challenge organizers. The training set CTCF peak calls were downloaded from the challenge website. All data from the challenge is used with permission from the organizers.

### Model architectures

We evaluated the performance of a variety of neural network models for our tasks. For models that used only epigenomic signal as input, we considered all models that had between 1 and 5 layers and all powers of 2 between 1 and 4096 neurons per layer.

For models that used only nucleotide sequence as input, we considered two different types of models. The first are fully dense networks similar to those that used only epigenomic signal. These models had between 1 and 3 layers with all powers of 2 between 1 and 1024 neurons per layer. The second are convolutional models that are composed of a variable number of convolutional layers followed by max pooling layers and ending with a single dense layer. These convolutional models had between 1 and 3 convolutional layers, between 1 and 256 filters per convolutional layer, and between 1 and 1024 nodes in the final dense layer. The convolutional layers used a kernel of size 8 and a stride of 1. The max pooling layers had a kernel of size 4 and a stride of 4.

The models that used both nucleotide sequence and epigenomic signal were composed of one of the nucleotide models above and one of the epigenomic models. The final hidden layers of the two models were concatenated together and fed through an additional hidden layer before the output. Rather than consider all potential model architectures that utilized nucleotide sequence, we limited our evaluation to only 100 randomly selected model architectures for computational reasons.

In all models, both the convolutional layers and the hidden dense layers used ReLU activations, where *f* (*x*) = *max*(0, *x*).

### Model training

The neural network models were trained in a standard fashion for neural network optimization. This involved using the Adam optimizer [23] and a binary cross-entropy loss. All model hyperparameters were set to their defaults as specified by Keras version 2.0.8 [24], and no additional regularization was used. The models were trained on balanced mini-batches of size 32, and an epoch was defined as 400 mini-batches. Training proceeded for 100 epochs, but was stopped early if performance on a balanced validation minibatch of size 3,200 did not improve after five consecutive epochs.

The gradient boosted decision tree models were trained using XGBoost [25]. The default values were used for all parameters, except that training progressed for 300 iterations, instead of 100, and the maximum depth of each tree was set to 6, instead of 3. The model was trained using a binary logistic loss and a L2 regularization strength of 1. A single model was trained for each input feature set. These models are then evaluated using the first *N* trees, using *N* between 1 and 300, to get the performance of models of varying complexity. Because subsampling is not used, this procedure is identical to independently training models of varying sizes.

The training, validation, and test sets consisted of different genomic loci depending on the model evaluation setting. In the cross-chromosomal setting, the validation set was derived from chromosome 2 and the test set was derived from chromosome 1 for both tasks. For the gene expression task, the training set consisted of all genes in chromosomes 3 through 22, while for the TAD boundary prediction task, it consisted of all 40 kbp bins in chromosome 3. In the cross-cell type setting, the training, validation, and test sets were derived from chromosomes 2 through 22 in the gene expression task and chromosomes 2 and 3 in the TAD boundary prediction task. In the hybrid setting, the training and validation sets were the same as in the cross-cell type setting, but the test set for both tasks were samples derived from chromosome 1. We chose to hold the training set constant between the cross-cell type and hybrid approaches, rather than the test set, in order to demonstrate that models trained on the same data exhibit markedly different trends with respect to model complexity depending on the evaluation set.

Depending on the evaluation setting, these models were also trained on either a single, or multiple, cell types. In all cases, models were evaluated on data derived from the H1 cell line (E003). In the cross-chromosomal setting, models for both tasks were also trained on data from the H1 cell line (E003). For the gene expression task in both other settings, samples drawn from spleen (E113), H1 BMP4 derived mesendoderm cultured cells (E004), CD4 memory primary cells (E037), and sigmoid colon (E106) were used as the validation set, and all other cell types (excluding the H1 cell line) were used as the training set (see Supplementary Table S1). For predicting TAD boundaries, the validation set was drawn from GM12878 (E116) and the training set consisted of all other cell lines (excluding the H1 cell line).

## Supplement

**Figure S1:**
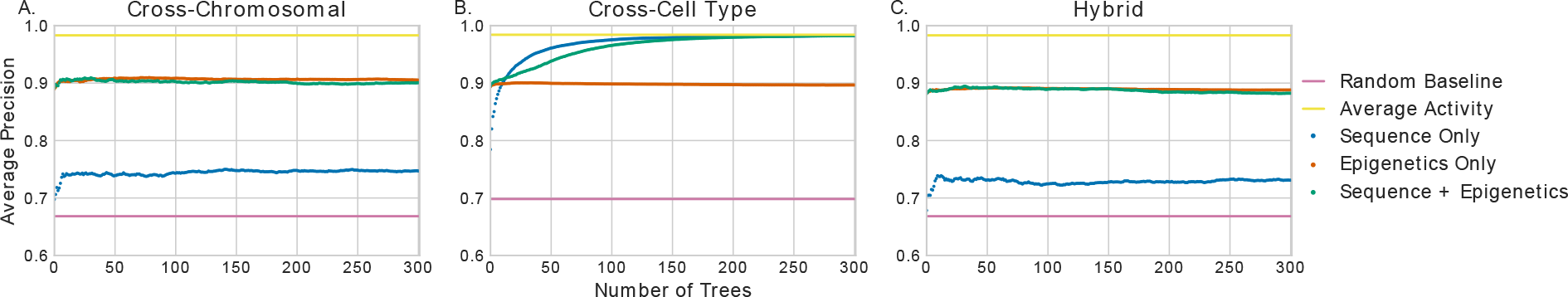
The performance of a gradient boosted decision tree classifier with a varying number of trees in three predictive settings for predicting gene expression. The figure plots the average precision (AP) of a gradient boosted decision tree model predicting gene expression as a function of model complexity. Evaluation is performed via (a) cross-chromosome, (b) cross-cell type, and (c) a combination of cross-chromosome and cross-cell type validation. In each panel, each point represents the test set performance of a single trained model.

**Figure S2:**
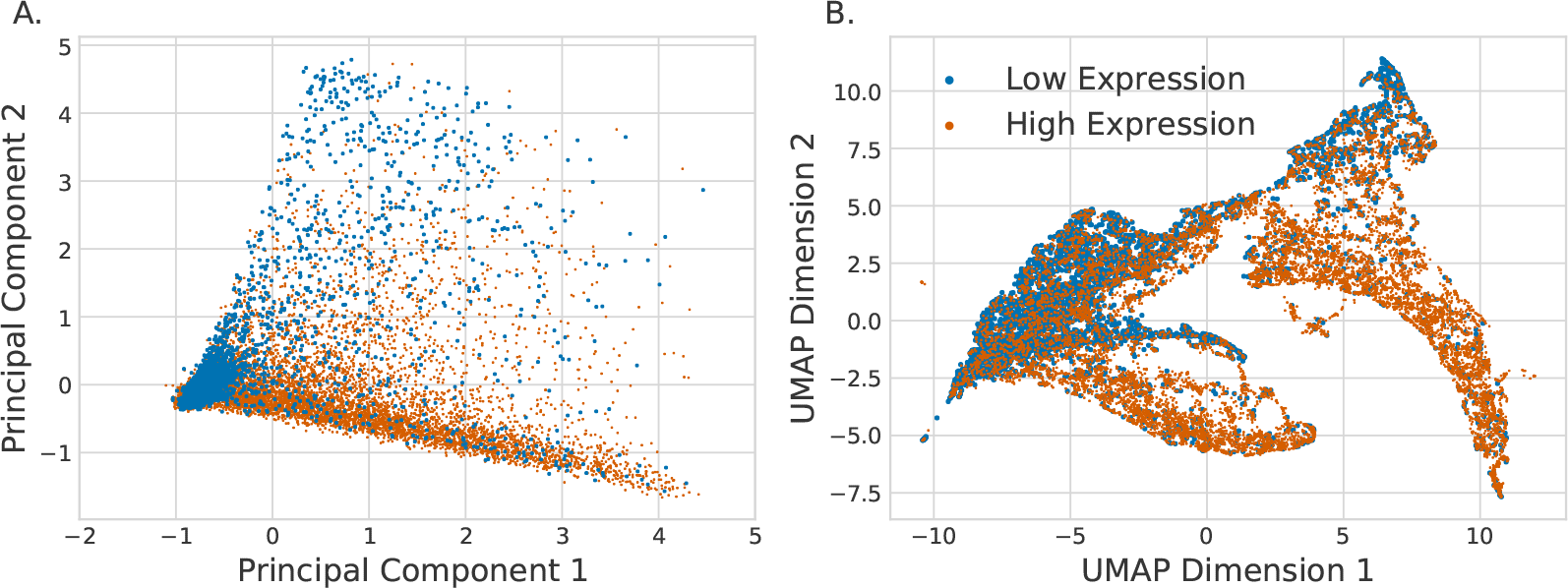
Projections of the epigenomic signal used to predict gene expression. The five histone modifications that were used to predict gene expression were projected down to two dimensions using (a) PCA and (b) UMAP. The projections are then colored by whether the gene is highly expressed (orange) or lowly expressed (blue) in H1.

**Table S1:** Training cell lines used to predict gene expression in the cross-cell type and hybrid evaluation settings. The Roadmap Epigenomics Mapping Consortium ID and biosample description are given for each.

| Roadmap ID | Biosample Summary |
| --- | --- |
| E004 | H1 BMP4 Derived Mesendoderm Cultured Cells |
| E005 | H1 BMP4 Derived Trophoblast Cultured Cells |
| E006 | H1 Derived Mesenchymal Stem Cells |
| E007 | H1 Derived Neuronal Progenitor Cultured Cells |
| E011 | hESC Derived CD184+ Endoderm Cultured Cells |
| E012 | hESC Derived CD56+ Ectoderm Cultured Cells |
| E013 | hESC Derived CD56+ Mesoderm Cultured Cells |
| E016 | HUES64 Cell Line |
| E024 | 4star |
| E027 | Breast Myoepithelial Cells |
| E028 | Breast vHMEC |
| E037 | CD4 Memory Primary Cells |
| E038 | CD4 Naive Primary Cells |
| E047 | CD8 Naive Primary Cells |
| E050 | Mobilized CD34 Primary Cells Female |
| E053 | Neurosphere Cultured Cells Cortex Derived |
| E054 | Neurosphere Cultured Cells Ganglionic Eminence Derived |
| E055 | Foreskin Fibroblast Primary Cells skin01 |
| E056 | Foreskin Fibroblast Primary Cells skin02 |
| E057 | Foreskin Keratinocyte Primary Cells skin02 |
| E058 | Foreskin Keratinocyte Primary Cells skin03 |
| E059 | Foreskin Melanocyte Primary Cells skin01 |
| E061 | Foreskin Melanocyte Primary Cells skin03 |
| E062 | Peripheral Blood Mononuclear Primary Cells |
| E065 | Aorta |
| E066 | Adult Liver |
| E070 | Brain Germinal Matrix |
| E071 | Brain Hippocampus Middle |
| E079 | Esophagus |
| E082 | Fetal Brain Female |
| E084 | Fetal Intestine Large |
| E085 | Fetal Intestine Small |
| E087 | Pancreatic Islets |
| E094 | Gastric |
| E095 | Left Ventricle |
| E096 | Lung |
| E097 | Ovary |
| E098 | Pancreas |
| E100 | Psoas Muscle |
| E104 | Right Atrium |
| E105 | Right Ventricle |
| E106 | Sigmoid Colon |
| E109 | Small Intestine |
| E112 | Thymus |
| E113 | Spleen |
| E114 | A549 EtOH 0.02pct Lung Carcinoma |
| E116 | GM12878 Lymphoblastoid |
| E117 | HeLa-S3 Cervical Carcinoma |
| E118 | HepG2 Hepatocellular Carcinoma |
| E119 | HMEC Mammary Epithelial |
| E120 | HSMM Skeletal Muscle Myoblasts |
| E122 | HUVEC Umbilical Vein Endothelial Cells |
| E123 | K562 Leukemia |
| E127 | NHEK-Epidermal Keratinocytes |
| E128 | NHLF Lung Fibroblasts |

**Figure S3:**
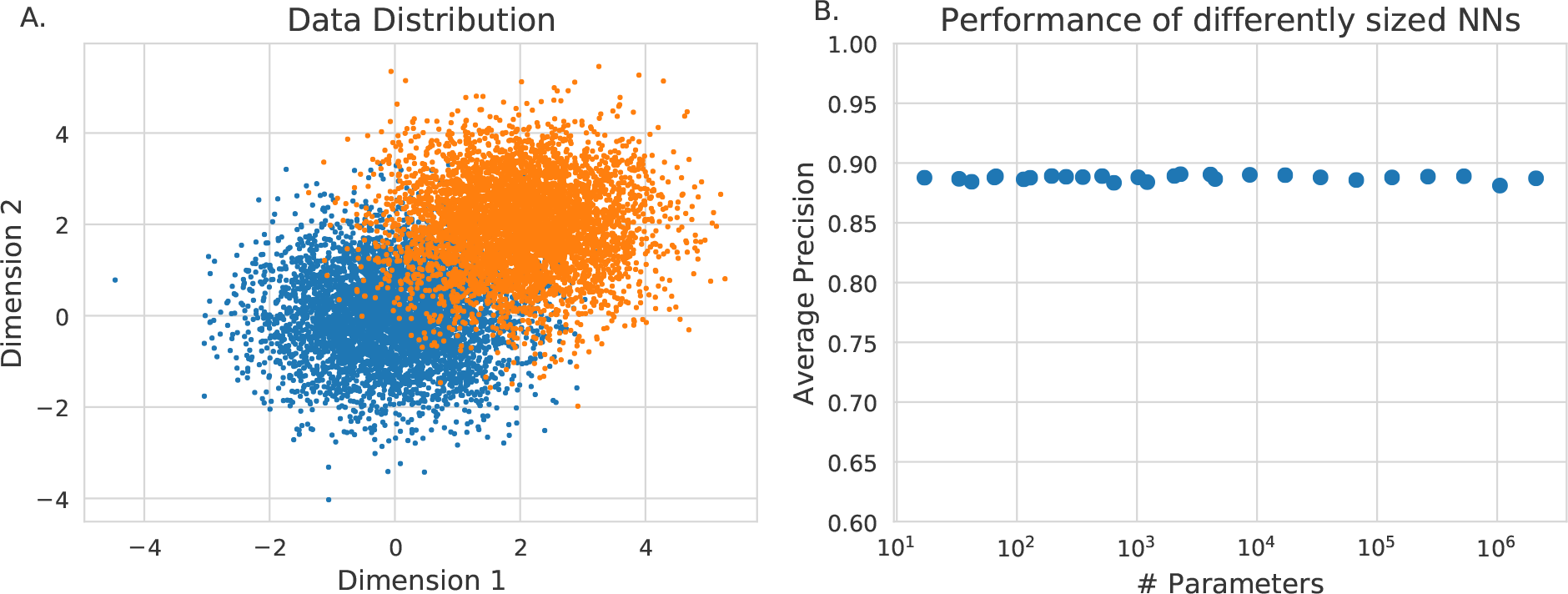
Classification performance of neural networks when the decision boundary is simple. (a) Random data was generated from two overlapping 2D Gaussian distributions. (b) Neural networks of increasing size were trained to classify points as either orange or blue and evaluated using the average precision. The y-axis is scaled to the same range as Figure 1d/e/f to demonstrate a similar trend.

**Figure S4:**
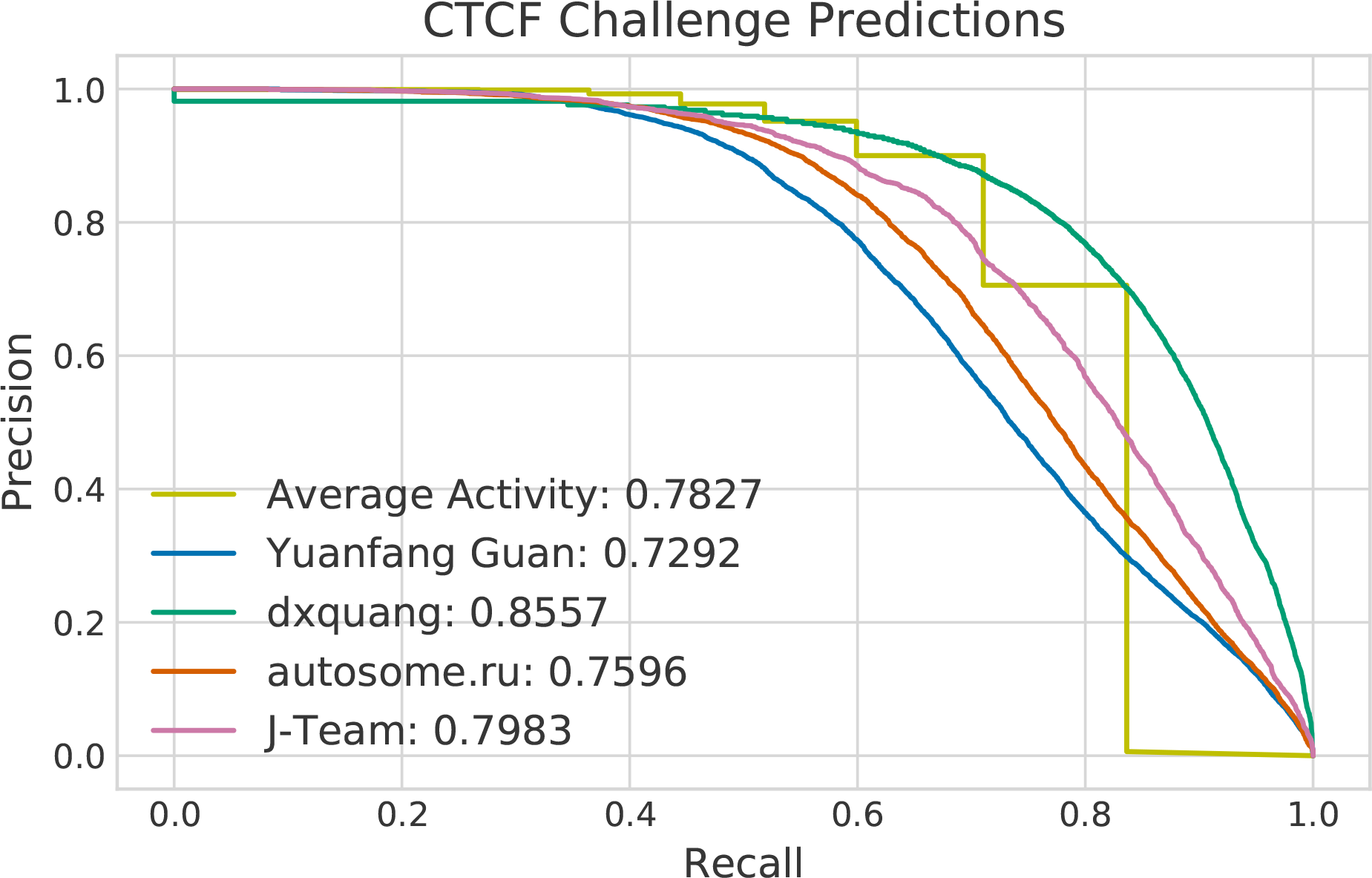
Performance of the top four participants in the ENCODE-DREAM TF binding challenge compared to the average activity baseline at predicting CTCF in iPSC. Precision-recall curves for each of the top four participants in the ENCODE-DREAM TF binding challenge, as well as a precision-recall curve for the average activity baseline at the same task. The average precision of each approach is shown in the legend.

**Figure S5:**
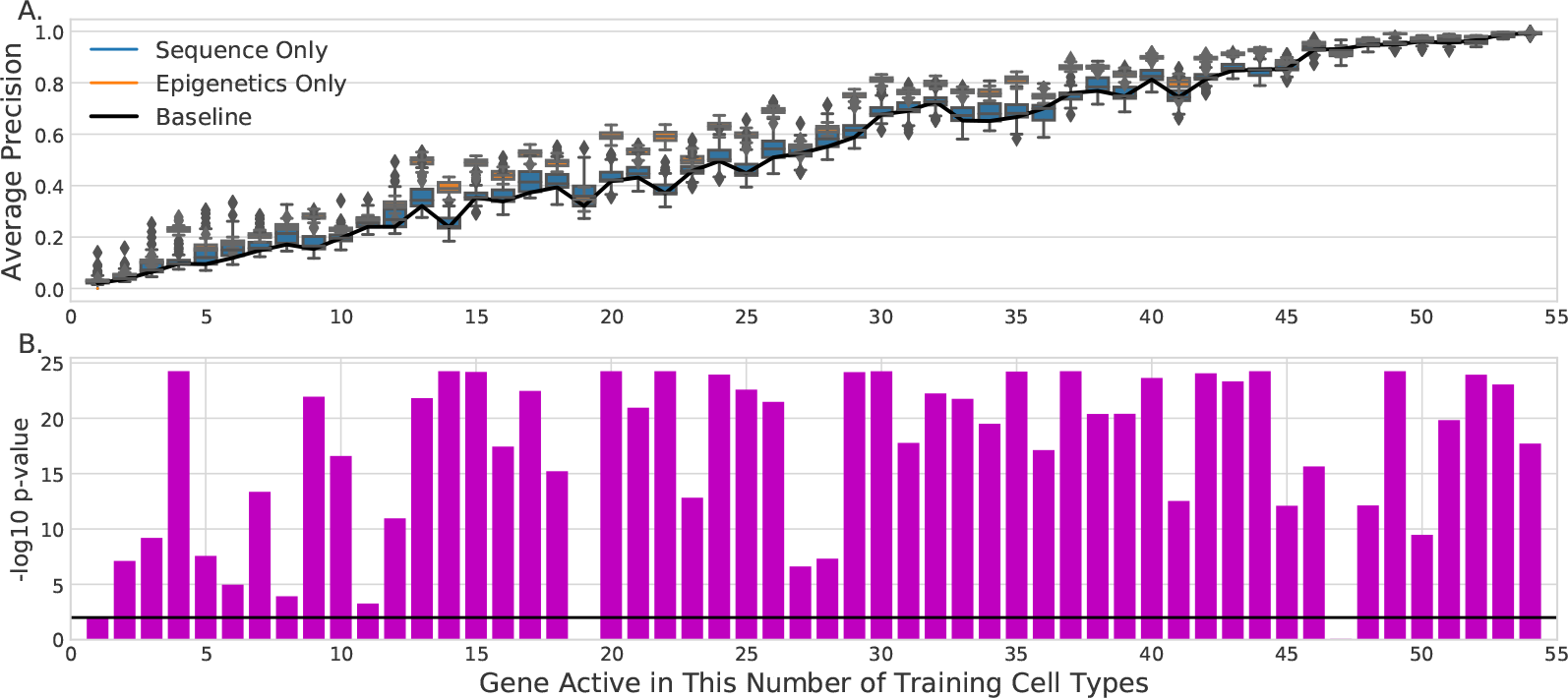
Epigenomic signal yields more predictive models than nucleotide sequence in the cross-cell type setting when locus-specific biases are factored out. Genes in the cross-cell type setting were split into 54 groups based on the number of training and validation set cell types that they are active in. (a) For each group, the AP score was calculated using the predicted probabilities from models that use only nucleotide sequence or use only epigenomic signal. Each box shows the three quartile values, with whiskers extending to 1.5 the inter-quartile range. (b) The AP scores from those two groups were then compared using a one-sided Mann-Whitney U test. The −log10 p-values of this test are displayed for each group. The null hypothesis is rejected for most groups, indicating that models that use epigenomic signal outperform those that use only nucleotide sequence when the average activity is factored out of the evaluation. As expected, the epigenetics-only case is relatively better as the uncertainty increases, corresponding to the middle of the plot above.

